## Supplemental Table - strain list for "Conserved NIMA kinases regulate multiple steps of endocytic trafficking"

| **Strain** | **Genotype** |
| --- | --- |
| PHX1714 | *syb1714 (rab-11.1) mScarlet::rab-11.1* |
| RT3533 | *pwSi7[phyp7::GFP::rab-7]* |
| RT3596 | *pw27[nekl-2::aid]; ieSi57[peft-3::mRuby::tir1]; pwSi7[phyp7::GFP::rab-7]* |
| RT3597 | *pw29[nekl-3::aid]; ieSi57[peft-3::mRuby::tir1]; pwSi7[phyp7::GFP::rab-7]* |
| RT3649 | *pw27[nekl-3::aid]; ieSi57[peft-3::mRuby::tir1]; pwls439[GFP::rab-5]* |
| RT3650 | pw29[nekl-3::aid]; ieSi57[peft-3::mRuby::tir1]; pwls439[GFP::rab-5] |
| RT3995 | *bzls166[pmec4::mCherry]; pwSi202[phyp7::aman-2::mNeonGreen]* |
| RT4000 | *pwSi207[phyp7::sma-6::GFP]* |
| RT4266 | *pwSi363[phyp7::mig-14::GFP]* |
| RT4271 | *pwSi368[phyp7::daf-4::GFP]* |
| RT4337 | *pw43[tgn-38::gfp]* |
| WY1193 | *nekl-2(fd100)[nekl-2::mNeonGreen::3xFlag]); nekl-3(fd106)[nekl-3:: mKate2::3xFlag]* |
| WY1716 | *pw29[nekl-3::aid]; ie57[peft-3::mRuby::tir-1]; pwSi207[phyp7::sma-6::GFP]* |
| WY1717 | *pw29[nekl-3::aid]; ie57[peft-3::mRuby::tir-1]; pwSi363[phyp7::mig-14::GFP]* |
| WY1719 | *pw29[nekl-3::aid]; ie57[peft-3::mRuby::tir-1]; pwSi368[phyp7::daf-4::GFP]* |
| WY1728 | *pw27[nekl-2::aid]; ie57[peft-3::mRuby::tir-1]; pwSi207[phyp7::sma-6::GFP]* |
| WY1736 | *pw27[nekl-2::aid]; ie57[peft-3::mRuby::tir-1]; pw43[tgn-38::GFP] line 1* |
| WY1741 | *pw27[nekl-2::aid); ie57[peft-3::mRuby::tir-1]; pwSi363[phyp7::mig-14::GFP] line 1* |
| WY1758 | *pw29[nekl-3::aid]; ie57[peft-3::mRuby::tir-1]; pw43[tgn-38::GFP] line 1* |
| WY1763 | *pw1543[GFP::rab-5]* |
| WY1838 | *syb1714 (mScarlet::rab-11.1); pw29[nekl-3::aid]; pwSi10[phyp-7::BFP::tir-1]* |
| WY1844 | *syb1714 (mScarlet::rab-11.1); pw27[nekl-2::aid]; pwSi10[phyp-7::BFP::tir-1]* |
| WY1917 | *pw27[nekl-2::aid]; ie57[peft-3::mRuby::tir-1]; pwSi202[phyp7::aman-2::mNeonGreen]* |
| WY1935 | *pw29[nekl-3::aid]; ie57[peft-3::mRuby::tir-1]; pwSi202[phyp7::aman-2::mNeonGreen]* |
| WY1941 | *nekl-3::mKate CRISPR (fd106); pwls439[GFP::rab-5]* |
| WY1943 | *nekl-3::mKate CRISPR (fd106); pwSi7[phyp7::GFP::rab-7]* |
| WY1944 | *nekl-2::mKate CRISPR (fd127); pwls439[GFP::rab-5]* |
| WY1945 | *nekl-2::mKate CRISPR (fd127); pwSi7[phyp7::GFP::rab-7]* |
| WY1998 | *pw29[nekl-3::aid); ie57[peft-3::mRuby::tir-1]; pw43[tgn-38::GFP]; cup-5(fd395)* |
| WY2000 | *pw27[nekl-2::aid); ie57[peft-3::mRuby::tir-1]; pw43[tgn-38::GFP]; cup-5 (fd397)* |
| WY1464 | *nekl-2::mNeonGreen CRISPR (fd100); pwSi36[phyp7::mScarlet::apa-2]* |
| WY1469 | *nekl-3::mNeonGreen CRISPR (fd118); pwSi36[phyp7::mScarlet::apa-2]* |

**Table S1. List of strains used in this study**
